# A quantitative tumor–wide analysis of morphological heterogeneity of colorectal adenocarcinoma

**DOI:** 10.1101/2024.04.10.588907

**Authors:** Mihnea P. Dragomir, Vlad Popovici, Simon Schallenberg, Martina Čarnogurská, David Horst, Rudolf Nenutil, Fred Bosman, Eva Budinská

## Abstract

Morphologic heterogeneity of colorectal adenocarcinoma (CRC) is poorly understood. Previously, we identified morphological patterns associated with CRC molecular subtypes, and showed that these patterns have distinct molecular motifs (***Budinská et al., 2023***). Here, we evaluated the heterogeneity of these patterns across CRC. Three pathologists evaluated dominant, secondary, and tertiary morphology on four different tissue blocks per tumor in a pilot set of 22 CRCs (n=88). An artificial intelligence (AI) image analysis tool was trained using the pathologist-rated tumors to assess the morphologic heterogeneity on an expanded set of 161 CRCs (644 images). Heterogeneity was expressed as a combination of morphology patterns (morphotypes) across slides and normalized Shannon’s index (NSI). All pathologists agreed that the majority of tumors had 2-3 different dominant morphotypes, and that the complex tubular (CT) morphotype was the most common. AI analysis confirmed these observations in the full set. CT morphotype combined with all other dominant morphotypes within a tumor. Desmoplastic (DE) morphotype was rarely dominant and rarely combined with other dominant morphotypes. Mucinous (MU) was most often combined with solid/trabecular (TB) and papillary (PP). Most tumors showed medium or high NSI, but without clinical consequence. The proportion of DE morphotype was associated with higher T-stage, N-stage, metastasis, AJCC-stage, and shorter relapse-free survival, and MU morphotype was associated with higher grade, right side, microsatellite instability, and shorter overall survival. In conclusion, we observed high intratumoral morphological heterogeneity of CRC, and that not heterogeneity *per se*, but the proportion of certain morphotypes showed associations with clinical outcome. This has implications for molecular profiling of CRC.

## Introduction

In recent years, we have witnessed important efforts to classify colorectal cancer using molecular approaches. An initial attempt classified colorectal adenocarcinoma (CRC) in three subtypes: hypermutated (13%, molecular inactivation of DNA mismatch repair), ultramutated (2 − 3% POLE or POLD1 inactivation), and chromosomal instability (CIN) (85% somatic alterations) (***Cancer Genome Atlas Network, 2012***). This initial classification left us with a large subclass of CIN adenocarcinomas, and further classification attempts followed. We subsequently used transcriptomic data to classify CRC in 5 major subclasses: surface crypt-like, lower crypt-like, CIMP-H-like, mesenchymal/stromal and mixed (***Budinská et al., 2013***). These subclasses matched to a large extent well-known morphological patterns. Efforts to harmonize the multiple emerging molecular taxonomies of CRC led to the development of a transcriptome-based Consensus Molecular Subtype (CSM) classification (***Guinney et al., 2015***) which defined four classes: CMS1 (matching to the TCGA’s hypermutated class), CMS2 (characterized by high somatic copy number alterations a nd WNT and MYC activation), CMS3 (low somatic copy number alterations, KRAS mutations and metabolic deregulation), and CMS4 (high somatic copy number alterations, stromal infiltration, high levels of angiogenesis, and TGF−*β* activation) (***Guinney et al., 2015***).

Our group (***Popovici et al., 2017***) and later others (***Sirinukunwattana et al., 2021***) managed to replicate the transcriptome-based molecular classification of CRC using automatic analysis of readily available high-resolution digital haematoxylin-eosin (H&E) slides. These results confirmed the close relationship between tumor morphology and coding transcriptome and were further strengthen by our recent analysis of the molecular programs associated with specific morphological patterns (***Budinská et al., 2023***).

Building on these results, the purpose of the present study was to investigate intratumoral heterogeneity in CRC by addressing the following questions: (1) What is the extent of intratumoral heterogeneity across a CRC tumor? (2) How do different morphologies associate?, and (3) What impact with respect to clinical variables and prognosis does morphological admixture have on CRC? To address these questions, three gastrointestinal pathologists (without having any prior consensus discussion) scored primary, secondary, and tertiary morphologies at four different sites (sections) per tumor. Because there were some discrepancies between pathologists and, in order to advance this analysis to a more quantitative and automated level, an artificial intelligence (AI)- based model was developed to detect and quantify the CRC morphologies. Finally, we studied the correlations between the morphology buildup and the clinical covariates.

## Results

### Inter-observer reproducibility of intratumoral morphological heterogeneity assessment

To analyze morphological heterogeneity in CRC and its associations with clinical variables and patient’s prognosis, we took a multi-step approach. Initially, a subset of 88 slides corresponding to 22 tumors were evaluated by three expert/clinical pathologists.

Comparing the annotations of the three pathologists regarding morphotype classification regardless of dominant, secondary, and tertiary morphology, the MU, DE, and TB showed lower inter-observer variability, while for CT, PP, and SS the inter-observer variability was higher (Supplemental Figure SF1). The most commonly present morphotype found by all pathologists was CT: 60%-85% of slides Figure 1A) and 77%-99.5% of all slides of a tumor (referred from now on as tumor), (Figure 1B). With respect to frequency across tumors, two pathologists agreed that the least common morphotype is TB (present in 9%-36% of tumors) (Figure 1B), while across evaluated slides, the pathologists’ evaluation of the least frequent morphotype differed (TB for pat1, DE for pat2 and SE for pat3, Figure 1A).

**Figure 1.**
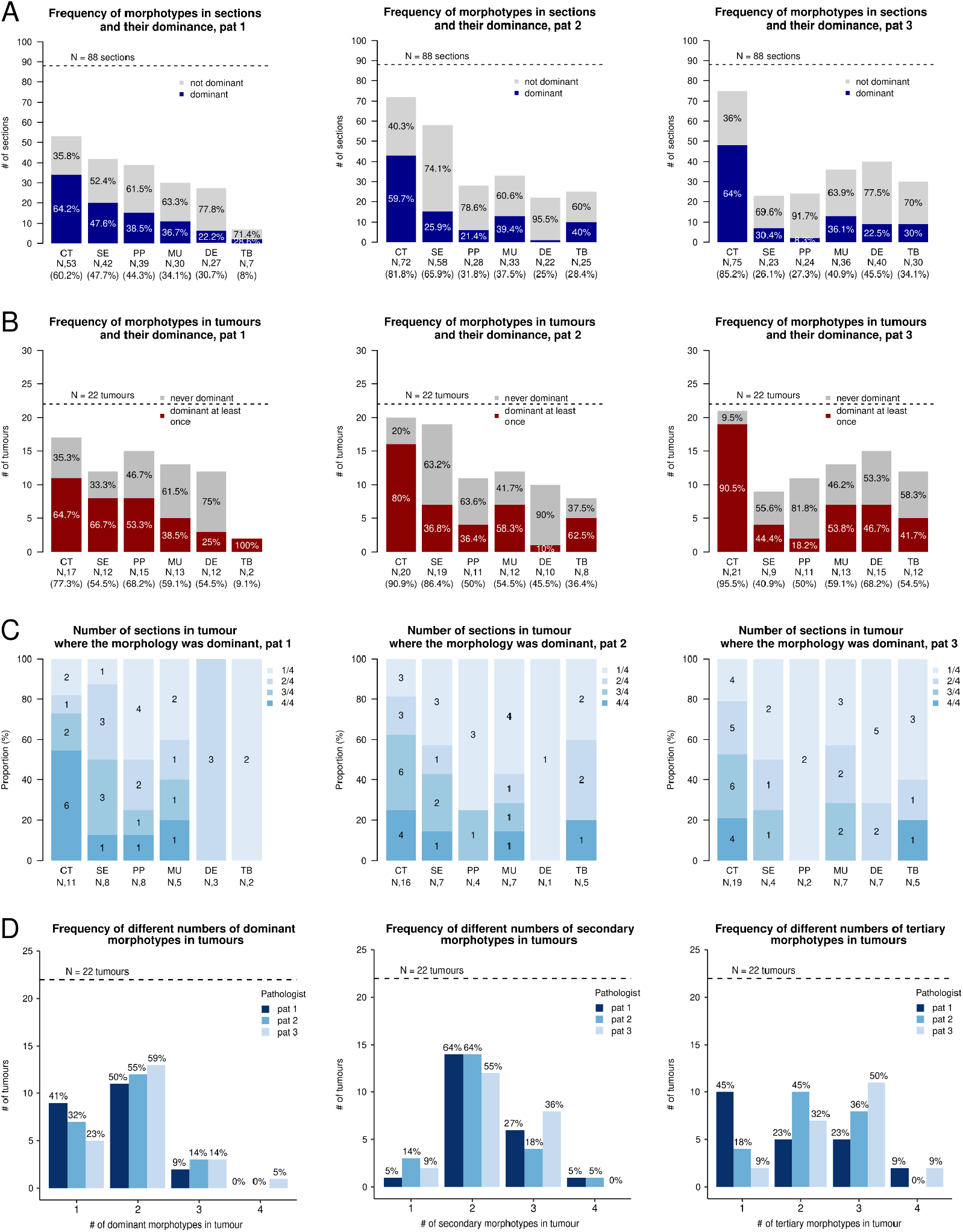
**A.** Frequency of morphotypes in sections with respect to their dominance. **B.** Frequency of morphotypes in tumors with respect to their dominance. **C.** Frequency of sections in tumors where the morphology was dominant. 1/4 means the morphotype was dominant in one of the four examined slides of the tumor etc. **D.** Frequency of different numbers of dominant (left), secondary (middle) and tertiary (right) morphotypes across the 4 examined sections, as evaluated by three expert pathologists.

If the CT morphotype was present in a tumor, all pathologists agreed that it was the dominant morphotype in at least one of the tumor’s slides, in most cases (ranging 65%-90.5%, Figure 1B). Additionally, DE morphology was never found dominant in all 4 examined sections of any tumor (Figure 1C). As much as 59%-73% of the tumors had two or three different morphotypes assigned as dominant across the four examined FFPE blocks. In one instance, one pathologist observed four different dominant morphotypes in a tumor (Figure 1D). For secondary morphotypes, this number was even larger, with 86%-95.0% tumors showing 2 to 4 different secondary morphotypes (Figure 1D). Tumors exhibited more than two different tertiary morphotypes in 55%-91% of cases (Figure 1D).

### Frequency, area, and dominance of the morphotypes as assessed by AI

The advantage of the AI-based image analysis was the automated quantification of percentage of area corresponding to each morphotype in the slides and the stability of the predictions, hence, we could extend our analysis to a more quantitative evaluation. Overall, 644 histopathology slides corresponding to 161 CRC were scored. Apart from the expected difference in tumor site (due to the design of the pilot study), no differences were observed between the full patient set and the pilot set evaluated by pathologists (Table 1).

**Table 1.** Distribution of clinical parameters of the individuals in the study, the full analysis and the pilot sets are compared. The pilot set is a subset of the full analysis set.

|  |  | full set | pilot set | p-value |
| --- | --- | --- | --- | --- |
| N |  | 161 | 22 |  |
| age (mean (SD)) |  | 66 (11) | 61 (14) | 0.055 |
| gender = M (%) |  | 89 (55.3) | 11 (50) | 0.812 |
| stage (%) |  |  |  | 0.093 |
|  | I | 24(14.9) | 0 (0.0) |  |
|  | II | 67 (41.6) | 10 (45.45) |  |
|  | III | 43 (26.7) | 10 (45.45) |  |
|  | IV | 27 (16.8) | 2 (9.1) |  |
| grade (%) |  |  |  | 0.170 |
|  | 1 | 19 (11.8) | 5 (22.7) |  |
|  | 2 | 95 (59.0) | 14 (63.7) |  |
|  | 3 | 47 (29.2) | 3 (13.6) |  |
| site (%) |  |  |  | 0.038 |
|  | right | 55 (34.2) | 11 (50.0) |  |
|  | transverse | 18 (11.2) | 0 (0.0) |  |
|  | left | 44 (27.3) | 10 (45.5) |  |
|  | rectosigmoid | 27 (16.8) | 1 (4.5) |  |
|  | rectum | 17 (10.6) | 0 (0.0) |  |
| pT stage (%) |  |  |  | 0.465 |
|  | T1 | 8 (5.0) | 1 (4.5) |  |
|  | T2 | 25 (15.5) | 1 (4.5) |  |
|  | T3 | 115 (71.4) | 19 (86.5) |  |
|  | T4 | 13 (8.1) | 1 (4.5) |  |
| pN stage (%) |  |  |  | 0.478 |
|  | N0 | 94 (58.4) | 10 (45.5) |  |
|  | N1 | 41 (25.5) | 8 (36.4) |  |
|  | N2 | 26 (16.1) | 4 (18.2) |  |
| pM stage = M1 (%) |  | 27 (16.8) | 2 (9.1) | 0.539 |
| MSI status = MSS (%) |  | 104 (79.4) | 20 (90.9) | 0.598 |

In the full set, the most commonly present morphotype in both individual sections (581/644, 90%) and all section of a tumor (158/161, 98%) was again CT, the least common morphotype was TB, found in 164/644 (25.5%) of slides and in 73/161 (45%) of tumors (Figure 2A and 2B). If present, CT morphotype occupied in median 50% tumor area within a section compared to other morphotypes, which occupied in median between 13% (SE) and 19% (PP) (Figure 2C). This correlates with the dominance of the morphotypes. If the CT morphotype was present in a tumor, it was the dominant morphotype in at least one of the tumor’s sections in most cases (123/158, 78%) (Figure 2B), compared to other morphotypes, which were not dominant most of the time (Figure 2B and 2D). Additionally, DE was never found dominant in all 4 examined sections of any tumor, most often was dominant in only 1 of 4 slides (15/23, 65%), similarly to SE (12/16, 75%), (Figure 2D). Overall, these initial observations match well with those of the expert pathologists.

**Figure 2.**
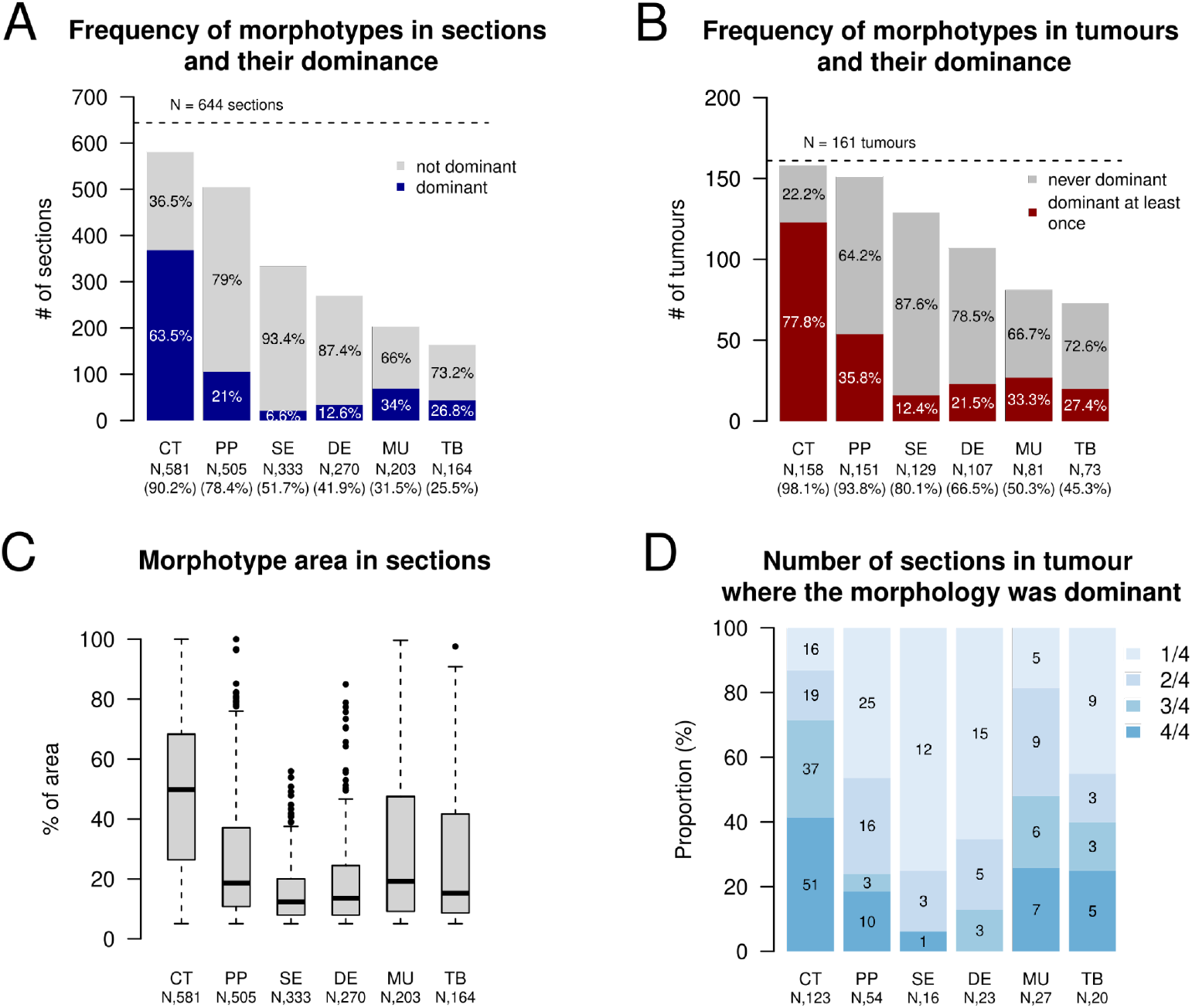
Frequency and area of dominant morphotypes across the examined sections and tumors. **A.** Frequency of morphotypes in sections with respect to their dominance. **B.** Frequency of morphotypes in tumors with respect to their dominance. **C.** Distribution of the fragment areas per individual morphotype, across all the sections. **D.** Frequency of sections in tumors where the morphology was dominant. 1/4 means the morphotype was dominant in one of the four examined slides of the tumor etc.

### Intratumoral morphological heterogeneity in CRC is variable

We further evaluated the intratumoral morphological heterogeneity from the perspective of morphotype dominance and by computing the normalized Shannon index (NSI) as a diversity measure on the proportions of the morphotypes across the four examined sections. As many as 54% (87/161) of the tumors had two or three different morphotypes assigned as dominant across the four examined H&E slides (we never observed four different dominant morphotypes in a tumor). For secondary and tertiary morphotypes, this number was even larger, with 86% (139/161) and 75% (120/161) tumors showing 2 to 4 different secondary or tertiary morphotypes, respectively.

Next, we examined the intratumoral patterns of dominant morphotype combinations (IPDMCs) and observed 39 groups representing four distribution patterns (4, 3+1, 2+2 and 2+1+1) (Figure 3A). 74/161 (46%) tumors only showed one dominant morphotype across the 4 examined blocks (51 CT, 10 PP, 7 MU, 5 TB and 1 SE), 52/161 (32%) and 20/161 (12%) tumors showed 2 dominant patterns distributed between 3+1 sections, and 2+2 sections, respectively. In addition, 15/161 (9%) tumors exhibited three dominant patterns distributed between 2+1+1 sections. The most frequent (>5) IPDMCs were 3xCT+1xPP (N=16 tumors), followed by 3xCT+1xDE (N=13), 2xCT+2xPP (N=8), 3xCT + 1xSE (N=6).

**Figure 3.**
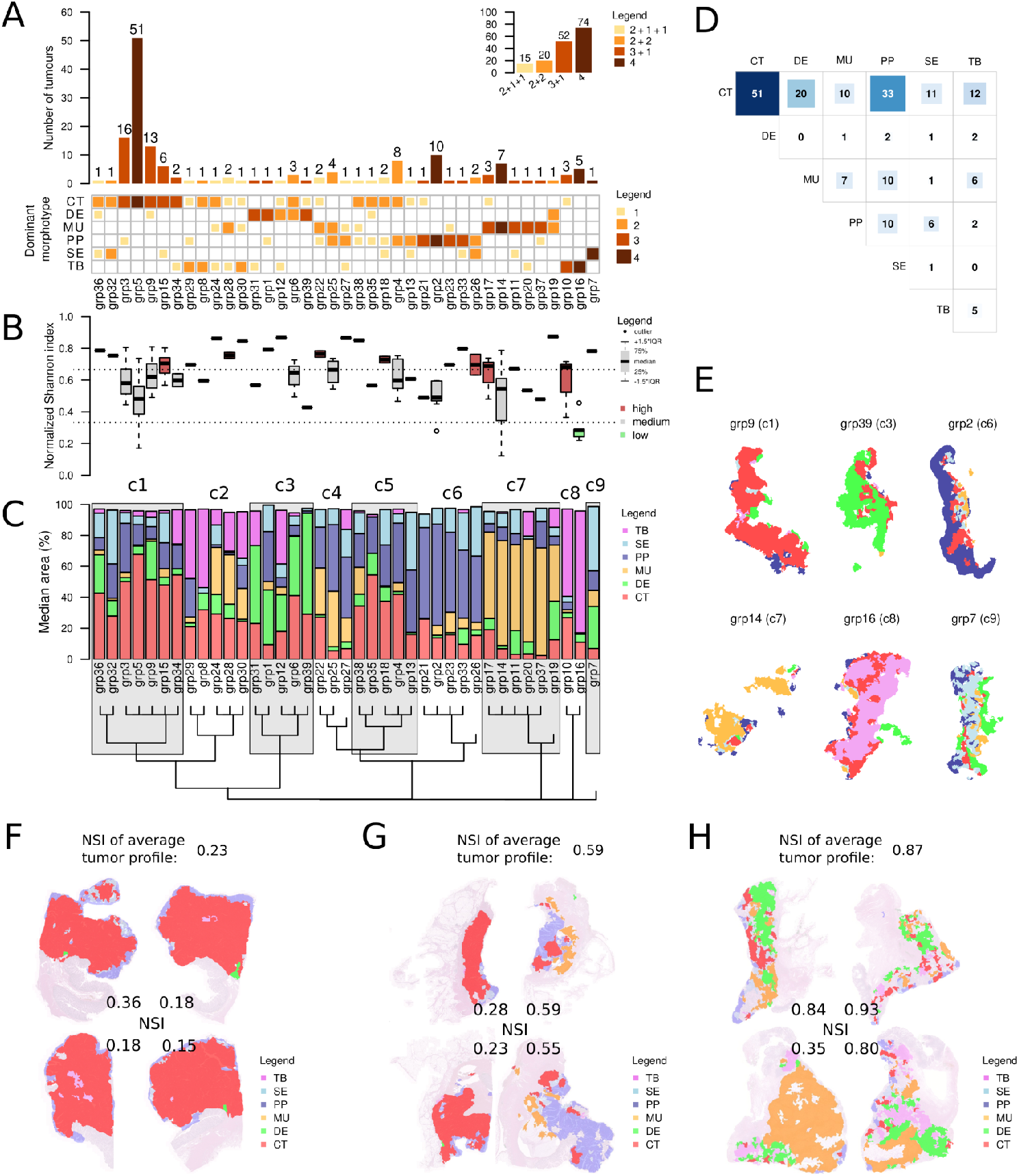
**A.** Observed intratumoral patterns of dominant morphotype combinations (IPDMCs) and their frequency (main barplot) and frequency of their distribution patterns (embedded top right barplot). **B.** Distribution of normalized Shannon index of median tumor profiles in the IPDMCs. **C.** Median morphotype area in the IPDMCs and their further clustering into 9 clusters. **D.** Frequencies of pairwise combinations of dominant morphotypes in the sections. **E.** Examples of representative tumor morphological areas of slides from selected IPDMCs clusters as identified by image analysis. **F-H.** Examples of intratumoral morphological heterogeneity as assigned by image analysis over four examined slides/blocks. Values of normalized Shannon index (NSI) of each slide and the average tumor profile are shown. **F.** Tumor with low heterogeneity across all slides, expressing one dominant morphotype (CT). **G.** Tumor with low heterogeneity in two slides and medium heterogeneity in two slides, expressing two dominant morphotypes (CT and PP) **H.** Tumor with high heterogeneity in all four sections, expressing two dominant morphotypes (DE and MU).

From the pairwise perspective, all the dominant morphotypes most often and almost uniquely combined with CT dominant morphotype, except for MU, which combined equally with PP and CT (10 cases each, Figure 3D). The most common pair was CT + PP (33 cases), followed by CT + DE (20 cases). We observed no combination of TB+SE and very few times a combination of MU+DE (1), MU+SE (1), DE+SE (1), DE+PP (2) and TB+DE (2), TB+PP (2).

For each IPDMCs group, we calculated its average morphotype composition based on which we computed the normalized Shannon index (NSI). This index is a measure of diversity that defines the internal heterogeneity of the sample from the perspective of all the morphotypes present and their fractions (Figure 3B). Interestingly, most tumors exhibited medium (NSI between 1/3 and 2/3) or high (NSI > 2/3) diversity and a few tumors exhibiting low diversity (NSI < 1/3, grp16, Figure 3B) were composed mostly of solid/trabecular morphotype. Hierarchical clustering of the average morphotype composition further divided the IPDMCs into 9 clusters (Figure 3C). Example sections of the IPDMCs can be found in Figure 3E. In Figure 3F-H we show examples of all 4 sections from tumors with low, medium and high morphological heterogeneity with their respective NSIs.

### Intratumoral morphotype heterogeneity and clinical parameters

We also looked for clinical and pathological parameters that associate with the morphotypes (expressed as proportion of the morphotype area in the tumor) and the degree of heterogeneity (expressed as the average NSI and as distribution patterns of the IPDMCs).

No significant associations were found between average NSI and the clinical parameters, or when testing the distribution patterns of IPDMCs (4, 3+1, 2+2 and 2+1+1) across stages (I-IV, Pearson’s *χ*^2^-test, p = 0.4561) or tumor site (left vs right, Pearson’s *χ*^2^-test, p=0.151).

The proportion of the DE morphotype increased and the proportion of PP decreased with increasing T-stage, N-stage, AJCC-stage and the presence of synchronous distant metastases (Figure 4A, Supplemental Tables ST1-ST8). The proportion of MU or TB morphotypes was associated with higher grade, right side and MSI. The proportion of CT morphotype was significantly higher in left-sided tumors and tumors of rectosigmoid and rectum. In addition, CT proportion was significantly lower in grade 3 and MSI tumors, similar to PP. Additionally, a larger area of TB morphotype was significantly associated with female gender. No significant associations were observed between SE morphotype and any of the clinical variables.

**Figure 4.**
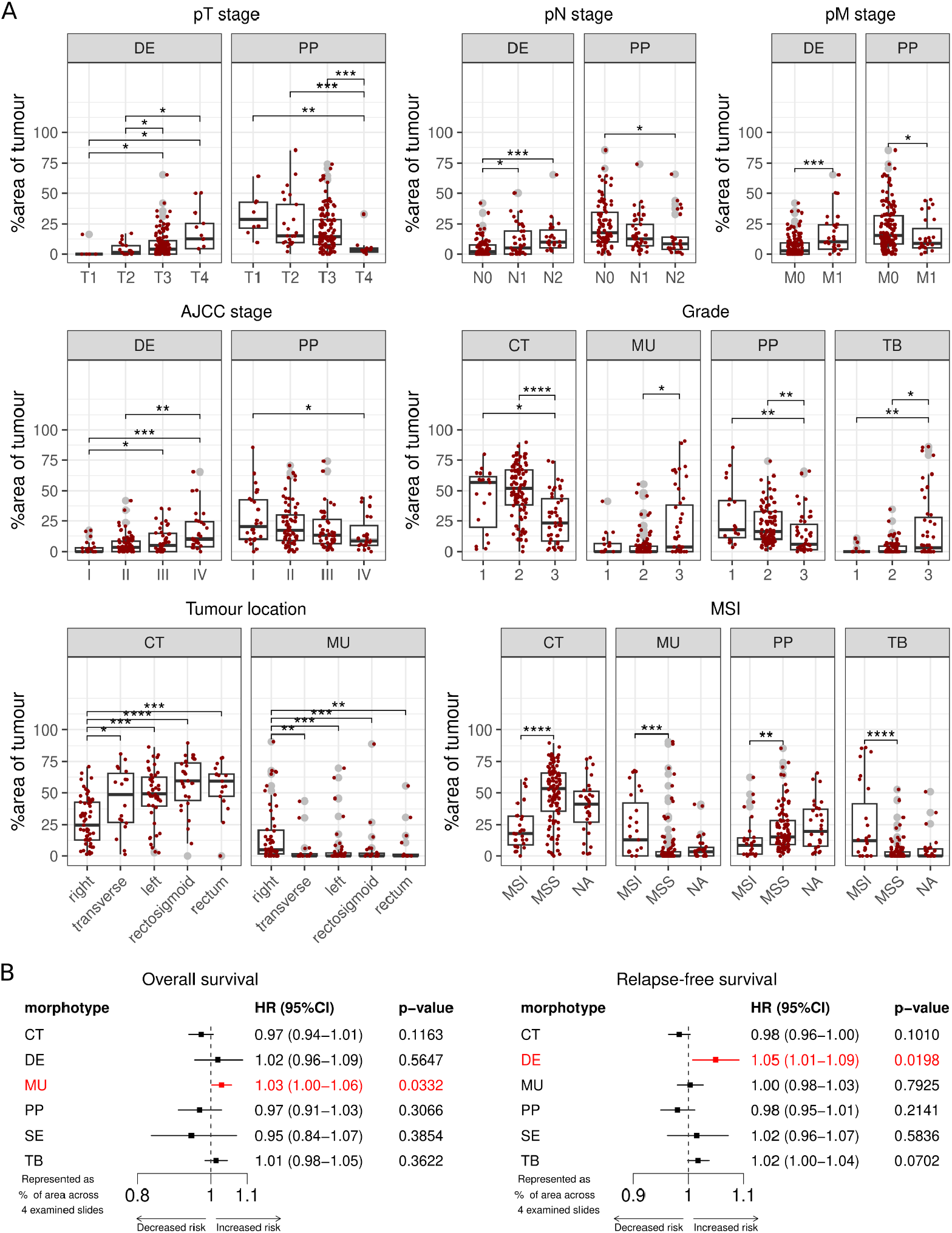
Associations between clinical variables and the morphotype area. **A.** Box plots of associations with categorical clinical variables at all stages. Middle line represents median, box represents interquartile range (IQR), whiskers represent ±1.5×IQR, and the gray dots represent outliers. Red dots represent single values (per tumor). Significant associations between the respective categories are indicated by connected lines and marked as follows: *: p<0.05; **: p<0.01; ***: p<0.001; ****: p<0.0001 (Mann-Whitney U-test between pairs of categories, if Kruskal-Wallis test significant). Additional associations and full statistics are provided in Supplemental Tables ST1-ST8. **B.** Forest plots of the association between overall survival (left) and relapse-free survival (right) and the morphotype area, calculated by Cox proportional hazards model, in stage I-III patients. Significant associations are highlighted in red.

Finally, we used the Cox proportional hazards model to model the association between the area of individual morphotypes (proportion across 4 examined slides) as well as NSI and overall survival (OS) and relapse-free survival (RFS) in stage I-III CRC patients. A 10% increase in the total area of the mucinous morphotype was associated with an approximately 10% increased risk of death in stage I-III CRC patients (Figure 4B left). A 10% increase in the total area of the desmoplastic morphotype is associated with an approximately 11% increased risk of recurrence (Figure 4B right). No significant association was found between NSI and survival. Due to the small sample size, we did not perform a survival analysis between the IPMDC groups.

## Discussion

In the present study we report that CRC is characterized by a high degree of morphological heterogeneity. Morphological heterogeneity in CRC is well recognized but insufficiently considered in diagnostic practice (***Remo et al., 2019***). Bulk tumor CRC transcriptome investigations culminated in the establishment of consensus molecular subtypes (CMS) that proved to have prognostic and predictive implications (***Guinney et al., 2015***). The original results on CMS already suggested that about 13% of the cases exhibited mixed features; we hypothesized that this could be caused by intratumoral heterogeneity. Indeed, the image-based CMS classification revealed that there is a high degree of intratumoral heterogeneity which is not accounted for during analysis of bulk sequencing data (***Popovici et al., 2017***; ***Sirinukunwattana et al., 2021***). However, even prior to the CMS, our research indicated that the CRC transcriptome subtypes are linked to various morphological patterns (***Budinská et al., 2013***). Having those patterns as a starting point, we later identified pattern-specific molecular programs in CRC (***Budinská et al., 2023***). Hence, we wanted to analyze the whole spectrum of morphological heterogeneity by using both expert pathologist and AI-based methodology and by considering four slides per tumor (from different FFPE blocks). We hypothesized that such an approach would provide a more comprehensive insight in morphologic heterogeneity of CRC, of potential clinical significance and providing essential cues for tumor sampling for molecular research.

We demonstrate that CRC is morphologically heterogeneous by analyzing 4 slides of 22 tumors (n = 88 slides) and integrating the assessments of three expert pathologists without prior consensus discussion. We observe that all pathologists discovered several morphotypes per slide and morphological differences between the slides of a single tumor. As expected (***Whitney-Miller, 2020***), there were some discrepancies between pathologists. We observed two elements that impacted interobserver variability: 1. pathologist experience (often the assessments of the two most experienced pathologists showed higher correlations for DE and MU morphology) and 2. interpathologist coordination (correlations between the assessment of the two pathologists from the same institution were higher than those with the third pathologist from a different institution). These discrepancies further outline the necessity for developing an objective and quantifiable AI tool that can automatically score CRC morphologies.

Therefore, we next confirmed the morphological heterogeneity of CRC by developing and using an AI deep learning model based on the annotations of the expert pathologists on an extended cohort of 161 CRCs (n = 644 slides). By visual morphological assessment and confirmed by AI-guided analysis we observed that >50% of cases harbor more than two dominant morphotypes on different sections from the same tumor, this number being even larger for secondary and tertiary morphotypes. This is reflected in the medium to high diversity of the normalized Shannon index, which combines proportions of different morphotypes regardless of dominance. Interestingly, the few tumors exhibiting low diversity were mostly of TB morphotype. We portend that the TB morphotype, with its complete loss of glandular architecture, may represent a final dedifferentiation stage, without a relation to any specific oncogenic program. Similarly, we found the CT morphotype to be more common than others, most often dominant and in combination with the other morphotypes. CT corresponds to WHO adenocarcinoma not-otherwise-specified (NOS), the most common subtype of CRC (***Shia et al., 2017***). CT might be considered as the most highly differentiated morphotype, most resembling normal colon mucosa architecture. Dedifferentiation of CT might then have TB as endpoint with other morphotypes as intermediates. In addition, SE morphotype may also be considered an original morphotype as reflecting the serrated neoplasia pathway, which is responsible for approximately 30% of cases of CRC (***O’Brien et al., 2015***).

Other common morphological combinations are MU with PP or MU with TB and PP with SE. These combinations could be partially explained by similar oncogenic programs, for example both the PP and MU morphologies are enriched in KRAS or BRAF driver mutations (***Remo et al., 2019***; ***Shia et al., 2017***; ***Patankar et al., 2018***; ***Kuroda et al., 2007***). DE combined almost exclusively with CT as the dominant morphotype. SE morphology very rarely combined with MU, which might be explained by its association with the serrated neoplasia pathway. Another potential explanation are morphological similarities, for example SE and PP are similar, and can be labeled by AI or pathologists differently on different slides, hence inducing artificial combinations.

Our observations have direct clinical relevance. Molecular analysis of CRC is routinely performed in the setting of recurrence and metastases, focusing on microsatellite status and driver mutations with direct implications for treatment (***Sveen et al., 2020***). Illustrative are the two cases reported by Büttner et al. in which the KRAS status differed between histotypes. In one case, mucinous morphology was *KRAS*-wild type, while non-mucinous morphology showed a *KRAS* G12D mutation. In a second case non-mucinous morphology showed a *KRAS* G12D mutation and a mucinous region a targetable *KRAS* G12C (***Büttner et al., 2018***). These observations indicate that in selecting the region for molecular analysis morphological heterogeneity needs to be taken into consideration and different morphotypes need to be included. AI-based image processing of histopathological slides as a guide for sample selection, as developed in this study, might become essential.

The correlations we found between morphotypes and clinicopathological parameters confirm well-known associations and offer a quantitative perspective to these associations. CT dominant morphotype associates with MSS (***Shia et al., 2017***), and MU with MSI and right side (***Green et al., 1993***). For the association between DE with higher T-stage and local and distant metastases the literature is more complex. DE can be further subcategorized into mature, intermediate and immature forms, the immature form having the worst prognosis (***Ueno et al., 2021***). Survival data showed that even a small proportion of MU morphotype (e.g. 10%) is indicative of shorter OS. In a previous study, it has been shown that WHO-defined mucinous CRCs (>50% mucinous morphology) have the worst prognosis, whereas there is no difference in survival between non-mucinous CRCs and CRCs with mucinous components (MU < 50%) (***Lee et al., 2013***). In addition, a large meta-analysis study involving more than 200,000 patients showed that mucinous differentiation (non-WHO defined) was associated with worse survival (HR 1.05 (1.02-1.08)) (***Verhulst et al., 2012***). These data highlight the need for a finer definition of the threshold, mainly using AI morphological quantification, to clarify the prognostic threshold of the MU morphotype. Our results challenge the WHO definition of mucinous CRC and support a 10% threshold. We further observed that the proportion of DE morphology associated with shorter RFS. The association between DE and survival is well defined and further stratified based by the type of desmoplasia (***Ueno et al., 2021***). Therefore, we suggest here that a quantitative approach is needed to gain a better prognostic understanding of the morphotypes.

Most molecular data on CRC are based upon samples disregarding intratumor morphological heterogeneity. Even studies performed on fresh or fresh-frozen tissue (***Cancer Genome Atlas Network, 2012***; ***ICGC/TCGA Pan-Cancer Analysis of Whole Genomes Consortium, 2020***) did not explicitly take morphological variants into consideration (***Shia et al., 2017***). AI-based analysis of H&E slides (***Minciuna et al., 2022***) or spatial transcriptomics (***Wang et al., 2023***; ***Wood et al., 2023***; ***Qi et al., 2022***) offer a much better insight in CRC heterogeneity. Indeed, such studies have further confirmed a very high degree of intratumoral heterogeneity and have begun to establish new links between morphology and the underlying molecular programs4,5,26. Our study reveals an additional point that will represent the bottleneck of CRC research: one slide/block is not enough. This puts the spotlight on the dissecting room of the pathology department and raises an essential practical question: what is the optimal sampling procedure for CRC, considering the morphologic heterogeneity, but also the definition of the tumor margin, which is essential if we want to understand the peritumoral immune infiltrate and the stromal response?

Mechanisms of morphological heterogeneity remain to be fully elucidated. They potentially involved are cancer cell genotype and transcriptome but also tumor microenvironment (including the stromal response, the immune infiltrate, and the microbiome). From this perspective, it would be of great interest to extend the histopathological characterization of tumor internal heterogeneity by inclusion of AI-based estimates of immune cell distribution within the tumor area and in close proximity.

Therefore, we can conclude by answering the questions raised in the beginnig: (1) intratumor morphologic heterogeneity in CRC is very high; (2) some morphological associations are more frequent while others are very rare, and this observation may shed light on the process of tumorigenesis and cancer clonality and; (3) the degree of heterogeneity is not associated with the clinical variables, on the contrary, it is the proportion of morphotypes that is associated with tumor stage, grade, location, and MSI.

Finally, we believe that AI-guided analysis of multiple H&E whole slides per single tumor is the most appropriate way to provide new insights into the associations between tumor (not only) morphological heterogeneity and clinical significance. Of course, more studies analyzing multiple distinct regions of a tumor are needed to draw definitive conclusions, but our data provide an important first step in this endeavor.

## Methods

### Study design and morphotype definition

The main research objective of the study was to analyze and quantify the morphological intratumoral heterogeneity in CRC and to understand the associations between different morphological patterns (morphotypes), and their impact on clinical parameters. *N* = 161 histologically-confirmed colorectal adenocarcinoma of stage I-IV were selected retrospectively from the hospital cohort of Masaryk Memorial Cancer Institute in Brno, Czech Republic, from cases that fit the selection criteria. The study was conducted in accordance with the principles of the Declaration of Helsinki and was approved by the ethical committee of Masaryk Memorial Cancer Institute. All patients provided written informed consent prior to enrollment.

Initial diagnostic H&E slide evaluation was performed by an expert pathologist according to the 2019 WHO classification (***Organisation mondiale de la santé, 2019***). For each tumor, we determined the T-stage, grade (G), localization, number of positive lymph nodes and total number of lymph nodes (N), and microsatellite instability (MSI) status. Patients with neoadjuvant treatment and patients with multiple tumors were excluded from the analysis.

For each case, we retrieved four distinct formalin-fixed paraffin-embedded (FFPE) blocks leading to a total of 644 blocks. The four selected FFPE blocks represented: i) the deepest invasion point at the serosa side, ii) the deepest invasion point at the insertion side of mesocolon/mesorectum, iii) transition point between the tumor and normal mucosa (including luminal side of the tumor), and iv) a representative tumor block. The tumor FFPE samples were cut into 5-micron thick consecutive slides and were stained with H&E. The H&E slides were scanned using the Pannoramic Midi (3DHISTECH) scanner at 20x magnification (0.234 microns/pixel resolution), with the same settings across all scans.

The study was then divided into the pilot phase and the AI-based phase. In the pilot phase of the study, three different pathologists (pat1, pat2, pat3) evaluated the morphologies in 4 slides from 22 selected tumors (88 slides). Since this set was limited in size, we matched the samples for age, gender, site and stage. Mainly left and right-sided tumors of stage II and III were considered for this subset (Table 1).

Morphotypes were defined by mapping frequently occurring morphologies of CRC as described in our previous publications (***Budinská et al., 2013***; ***Budinská et al., 2023***). These overlap in part with the carcinoma subtypes as defined in WHO 2019 (***Organisation mondiale de la santé, 2019***). We decided to use the term *morphotype* to indicate ‘pure morphology’, whereas the WHO’s *subtypes* are (inherently) morphologically heterogeneous. The six morphotypes were (see also Figure 1 in (***Budinská et al., 2023***)):

- complex tubular (CT), as found in WHO adenocarcinoma NOS (non-otherwise specified) and adenoma-like adenocarcinoma;
- solid/trabecular (TB), as found in WHO undifferentiated adenocarcinoma;
- mucinous (MU), as found in WHO mucinous adenocarcinoma, but without a 50% threshold;
- papillary (PA), mainly as found in WHO micropapillary adenocarcinoma;
- desmoplastic (DE), with no WHO histopathological equivalent but recognized as important tumor characteristic (***van Pelt et al., 2018***); and
- serrated (SE), as found in WHO serrated adenocarcinoma.

The pathologists evaluated the presence of the 6 different morphotypes and for each slide up to three different morphologies were scored in terms of their prevalence (dominant/primary – most prevalent, secondary, and tertiary). Of note, the morphotype choice was not made in consensus between the three pathologists, each pathologist independently scored the morphotypes and no common training sessions were conducted to create ‘common ground’ in classifying. We used this pilot phase to define the interobserver variability and to define morphotype regions for which all pathologists agreed regarding the annotations (ground truth). As other morphological patterns encountered in WHO histopathological subtypes, such as signet-ring cell carcinoma, adenosquamous carcinoma, and carcinomas with sarcomatoid components, are rare, these were not considered in our morphotype definitions.

In the AI-based phase, a deep learning image analysis model was trained to automatically detect the six morphotypes, and applied to all images in our collection (644 images). The model was trained on a subset of the 22-tumor set using regions with high degree of agreement among the three pathologists.

### AI image analysis

HALO® Image Analysis Platform (version 3.6.4134, Indica Labs, Inc., Albuquerque, New Mexico) and HALO® AI software (version 3.6.4134, Indica Labs, Inc., Albuquerque, New Mexico) were used for training a deep learning model (DenseNet V2) to segment the regions representing the six morphological patterns of interest. Expert annotations of representative regions for the morphological patterns were used as positive examples (for the 6 categories). The model was trained on lower magnification images (equivalent to 1.25x) to predict seven categories: the 6 categories of interest and a seventh “other” category covering the rest of the possible input cases, including background, normal epithelial tissue, fat, muscle, and other tumor regions not falling within the 6 categories of interest. To minimize the number of required samples, we employed a transfer learning approach, in which an initial base model is fine-tuned to the problem at hand. For each of the six categories of interest, we had between 3000 and 5000 image patches in the training set and about 10000 patches for the “other” category. The training stopped once the agreement (at whole slide level) between annotations and predictions (defined as a modified “intersection-over-union” coefficient: 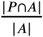, where *P* was the predicted region and *A* the annotated region), was greater than 0.9. for all categories of interest, on a set of independent images.

The AI model was applied to all images in the collection, which included 161 CRC, totaling 644 slides, also counting the initial 22 samples. The AI automatically segmented all samples and annotated them for the six morphotypes. The predicted regions corresponding to morphotypes (as produced by AI-based image analysis) were usually fragmented. To reduce the bias due to small regions that had higher chance of misclassification, we kept only the predicted regions covering at least 0.05 (i.e., ≥ 5%) of the tumor region. The correlation between the proportions of different morphotypes and intratumoral heterogeneity with several clinical and pathological parameters was analyzed.

### Statistical analysis

To measure inter-rater reliability between the three pathologists, we calculated pairwise the Cohen’s Kappa coefficient for the observation of each of the morphotypes separately, regardless of their prevalence. To quantify the heterogeneity of the morphotypes within the examined slides and tumors, we computed Shannon index (standard measure of sample diversity, based on proportions of its parts), and standardized this index by dividing the observed values by the maximum possible value (1.79 for 6 uniformly distributed morphotypes), so that we obtain index between 0 and 1. We call this index the Normalized Shannon Index (NSI).

To assess the association between categorical variables (clinical data between morphotype categories or between the two cohorts), the Pearson’s *χ*^2^ test was employed. The differences of morphotype area between groups were assessed using Kruskal-Wallis test (for more than two categories) or Mann-Whitney U test (for 2 categories). Age was tested between the two cohorts using Student’s t-test.

Hierarchical clustering was performed on centered log-ratio (clr)-normalized proportions of intratumoral patterns of dominant morphotype combinations (IPDMCs) with Euclidean distance and Ward algorithm to further define larger clusters.

To assess the impact of the morphotype area on both relapse-free survival, overall survival and NSI, we employed the Cox Proportional Hazards model. Where appropriate, we corrected for multiple hypothesis testing by controlling FDR using Benjamini-Hochberg method. Results were considered significant at FDR≤ 10%.

All statistical analyses were performed using `R` version 4.3.2 (***R Core Team, 2022***). Visualizations were performed using `R` packages `ggplot2` (3.4.4), `ggstatsplot` (v.0.12.1), `corrplot` (v 0.92), `forestploter` (v.1.1.1).

## Data availability

The datasets used and/or analyzed during the current study are available from the corresponding author on reasonable request.

## Acknowledgments

Supported by the Ministry of Health of the Czech Republic, grant no. 19-03-00298. All rights reserved. This project was funded by the Czech Science Foundation (GAČR), project no. GA19-08646S. This work was supported by the Berlin Institute of Health, Clinician Scientist Program (to M.P.D.), and DKTK Berlin Young Investigator Grant 2022 (to M.P.D.). The authors thank the RECETOX Research Infrastructure (No LM2023069) financed by the Ministry of Education, Youth and Sports, for its supportive background. Computational resources were provided by the e-INFRA CZ project (ID:90254), supported by the Ministry of Education, Youth and Sports of the Czech Republic. The work was also supported from Programme JAC - project SALVAGE (CZ.02.01.01/00/22_008/0004644) financed by the MEYS – co-funded by the European Union.

## Ethics approval

The study was approved by the ethical committee of Masaryk Memorial Cancer Institute. Ethical code: Č.J. 2015/1721/MOU, JID: MOU 73 866. All patients gave written informed consent in accordance with the Declaration of Helsinki prior to participating in the study.

