## Supplemental material for "A quantitative tumor–wide analysis of morphological heterogeneity of colorectal adenocarcinoma"

### Supplemental figures

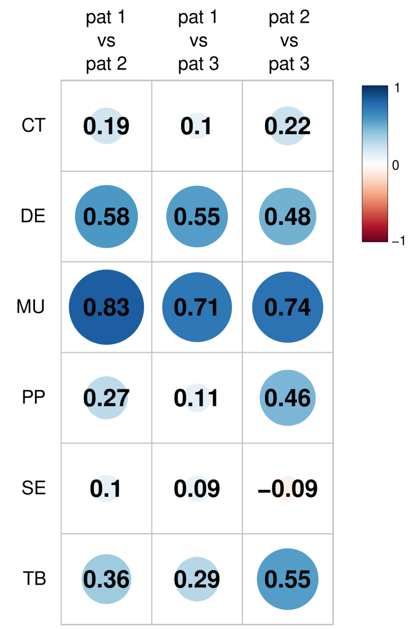


**Supplemental Figure SF1.** The pairwise analysis of operator agreement by Kappa coefficient for individual morphologies, where values represent *weak* $(0.4 - 0.59)$, *moderate* $(0.6 - 0.79)$, *strong* $(0.8 - 0.9)$, and almost perfect $(> 0.9)$ agreement.

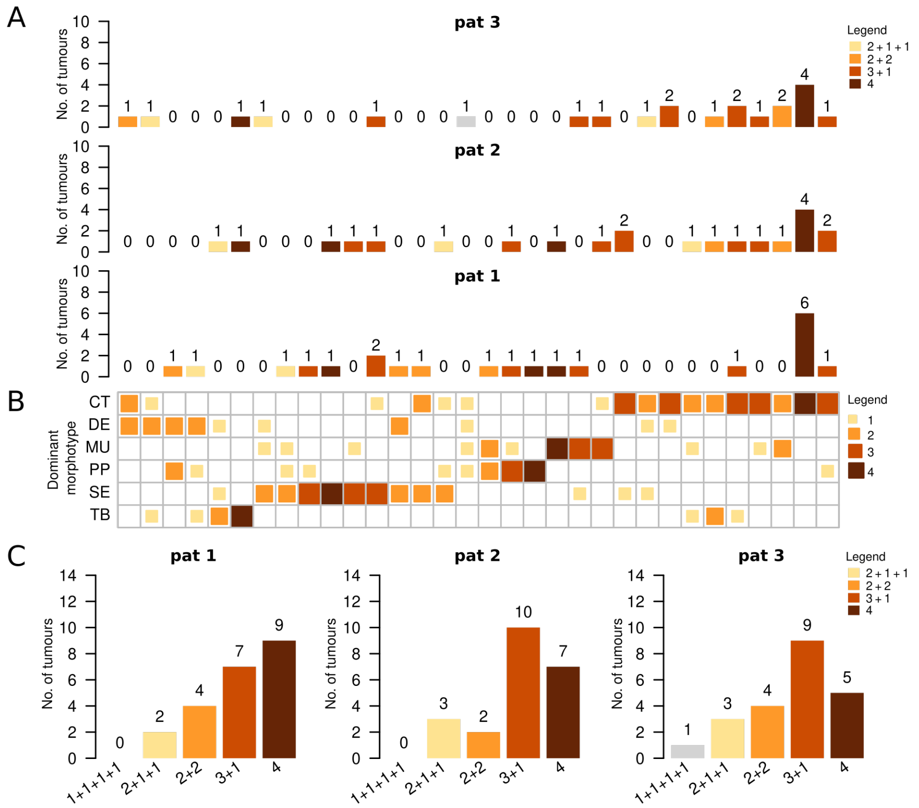

**Supplemental Figure SF2.** Observed intratumoural patterns of dominant morphotype combinations (IPDMCs) (B), their frequencies (A) and frequencies of their distribution patterns (C) as observed by three expert pathologists in the subset of 22 tumours.

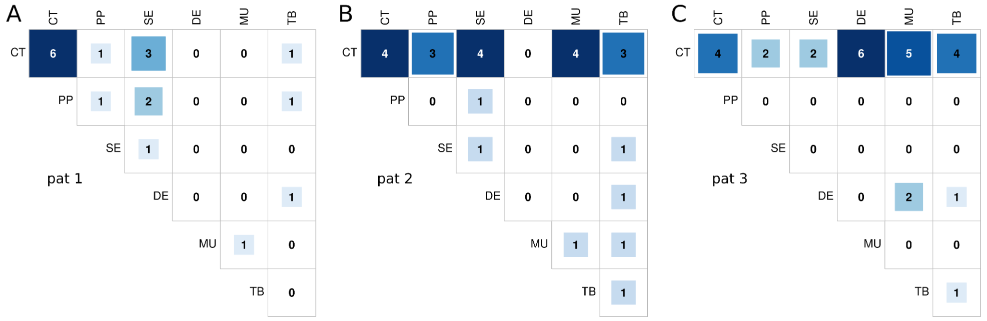

**Supplemental Figure SF3.** Frequencies of pairwise combinations of dominant morphotypes in the sections as evaluated by three pathologists. A. results for pat 1, B results for pat 2, C results for pat 3

### Supplemental tables

**Supplemental Table ST1.** Association between morphotype proportions and *gender* covariate.

| | F | M | p | p-adj |
|---|---|---|---|---|
| **n** | **72** | **89** | | |
| CT (median [IQR]) | 44.79 [18.03, 62.44] | 46.79 [26.75, 60.70] | 0.484 | 0.692 |
| DE (median [IQR]) | 5.53 [0.00, 12.69] | 2.60 [0.00, 9.30] | 0.069 | 0.144 |
| MU (median [IQR]) | 0.00 [0.00, 7.24] | 1.32 [0.00, 7.33] | 0.668 | 0.828 |
| PP (median [IQR]) | 12.63 [5.63, 26.66] | 16.87 [9.38, 32.85] | 0.075 | 0.145 |
| SE (median [IQR]) | 4.94 [1.80, 13.82] | 5.50 [1.75, 10.63] | 0.799 | 0.918 |
| **TB (median [IQR])** | **2.90 [0.00, 10.88]** | **0.00 [0.00, 2.74]** | **0.003** | **0.009** |

**Supplemental Table ST2.** Association between morphotype proportions and *AJCC stage* covariate.

| | I | II | III | IV | p | p-adj |
|---|---|---|---|---|---|---|
| **n** | **24** | **67** | **43** | **27** | | |
| CT (median [IQR]) | 46.90 [35.32, 65.42] | 44.84 [20.63, 61.90] | 50.68 [13.69, 59.29] | 43.00 [30.36, 60.02] | 0.929 | 0.949 |
| **DE (median [IQR])** | **0.00 [0.00, 3.09]** | **3.53 [0.00, 8.66]** | **5.10 [0.00, 14.63]** | **10.28 [4.08, 24.30]** | **<0.001** | **<0.001** |
| MU (median [IQR]) | 0.00 [0.00, 3.73] | 1.33 [0.00, 12.63] | 1.57 [0.00, 10.46] | 0.00 [0.00, 4.65] | 0.466 | 0.692 |
| PP (median [IQR]) | 20.30 [10.06, 42.55] | 17.19 [9.22, 29.87] | 13.10 [6.60, 26.21] | 8.87 [5.14, 21.14] | 0.046 | 0.110 |
| SE (median [IQR]) | 5.98 [1.63, 16.95] | 5.55 [2.31, 12.71] | 5.06 [1.85, 9.46] | 4.19 [0.00, 12.32] | 0.838 | 0.924 |
| TB (median [IQR]) | 0.65 [0.00, 5.04] | 0.00 [0.00, 6.62] | 0.00 [0.00, 7.19] | 1.29 [0.00, 5.33] | 0.996 | 0.996 |

**Supplemental Table ST3.** Association between morphotype proportions and *grade* covariate.

| | 1 | 2 | 3 | p | p-adj |
|---|---|---|---|---|---|
| **n** | **19** | **95** | **47** | | |
| **CT (median [IQR])** | **56.93 [19.71, 61.70]** | **52.01 [38.28, 66.80]** | **23.33 [8.82, 43.43]** | **<0.001** | **<0.001** |
| DE (median [IQR]) | 3.64 [0.00, 9.80] | 3.66 [0.00, 10.60] | 4.30 [0.00, 12.83] | 0.519 | 0.692 |
| **MU (median [IQR])** | **0.00 [0.00, 6.73]** | **0.00 [0.00, 4.42]** | **3.81 [0.00, 38.09]** | **0.026** | **0.073** |
| **PP (median [IQR])** | **17.66 [11.10, 41.91]** | **16.13 [10.10, 32.60]** | **6.14 [1.56, 22.13]** | **<0.001** | **0.002** |
| SE (median [IQR]) | 8.40 [4.95, 13.34] | 5.28 [1.80, 11.67] | 3.91 [0.00, 12.25] | 0.060 | 0.132 |
| **TB (median [IQR])** | **0.00 [0.00, 0.00]** | **0.00 [0.00, 4.49]** | **3.20 [0.00, 27.86]** | **0.001** | **0.006** |

**Supplemental Table ST4.** Association between morphotype proportions and *tumor site* covariate.

| | right | transverse | left | rectosigmoid | rectum | p | p-adj |
|---|---|---|---|---|---|---|---|
| **n** | **55** | **18** | **44** | **27** | **17** | | |
| **CT (median [IQR])** | **24.62 [12.69, 42.92]** | **48.56 [26.82, 65.46]** | **49.16 [39.79, 62.50]** | **59.51 [44.31, 73.78]** | **59.32 [47.16, 64.91]** | **<0.001** | **<0.001** |
| DE (median [IQR]) | 4.73 [1.43, 14.31] | 6.69 [0.53, 9.11] | 1.49 [0.00, 8.26] | 1.69 [0.00, 8.54] | 3.66 [0.00, 14.01] | 0.178 | 0.329 |
| **MU (median [IQR])** | **4.94 [1.69, 20.55]** | **0.00 [0.00, 1.79]** | **0.00 [0.00, 2.43]** | **0.00 [0.00, 2.11]** | **0.00 [0.00, 1.32]** | **<0.001** | **<0.001** |
| PP (median [IQR]) | 12.51 [4.80, 32.88] | 12.58 [4.07, 30.99] | 19.74 [9.93, 34.19] | 12.39 [7.86, 19.52] | 15.17 [13.10, 23.44] | 0.379 | 0.674 |
| SE (median [IQR]) | 5.74 [1.65, 13.97] | 4.16 [1.39, 11.80] | 6.02 [1.98, 13.02] | 5.28 [2.84, 12.00] | 2.72 [0.00, 10.63] | 0.858 | 0.924 |
| **TB (median [IQR])** | **1.30 [0.00, 13.59]** | **4.22 [0.00, 9.95]** | **0.00 [0.00, 4.49]** | **0.00 [0.00, 3.20]** | **1.29 [0.00, 3.51]** | **0.038** | **0.096** |

**Supplemental Table ST5.** Association between morphotype proportions and *pathologic T stage (pT)* covariate.

| | **T1** | **T2** | **T3** | **T4** | **p** | **p-adj** |
|---|---|---|---|---|---|---|
| **n** | **8** | **25** | **115** | **13** | | |
| CT (median [IQR]) | 47.81 [45.98, 53.60] | 47.36 [32.29, 66.98] | 44.75 [18.84, 60.67] | 52.01 [29.38, 66.62] | 0.578 | 0.750 |
| **DE (median [IQR])** | **0.00 [0.00, 0.00]** | **1.39 [0.00, 7.01]** | **4.11 [0.00, 11.18]** | **12.67 [4.46, 25.44]** | **0.001** | **0.005** |
| MU (median [IQR]) | 0.00 [0.00, 0.00] | 1.57 [0.00, 4.83] | 1.49 [0.00, 9.67] | 0.00 [0.00, 10.13] | 0.059 | 0.132 |
| **PP (median [IQR])** | **28.81 [21.54, 42.55]** | **15.17 [9.53, 40.71]** | **14.47 [7.90, 28.55]** | **3.48 [1.66, 5.00]** | **<0.001** | **<0.001** |
| SE (median [IQR]) | 2.41 [1.21, 15.97] | 8.13 [2.03, 15.23] | 5.55 [1.87, 12.71] | 1.90 [0.00, 3.65] | 0.073 | 0.145 |
| TB (median [IQR]) | 0.00 [0.00, 4.62] | 0.00 [0.00, 4.91] | 0.00 [0.00, 6.54] | 3.12 [0.00, 6.69] | 0.803 | 0.918 |

**Supplemental Table ST6.** Association between morphotype proportions and *pathologic N stage (pN)* covariate.

| | **N0** | **N1** | **N2** | **p** | **p-adj** |
|---|---|---|---|---|---|
| **n** | **94** | **41** | **26** | | |
| CT (median [IQR]) | 45.80 [21.50, 63.39] | 47.38 [27.79, 59.26] | 40.31 [10.68, 59.46] | 0.474 | 0.692 |
| **DE (median [IQR])** | **2.02 [0.00, 7.63]** | **5.10 [0.00, 19.09]** | **9.90 [5.15, 19.99]** | **<0.001** | **<0.001** |
| MU (median [IQR]) | 0.00 [0.00, 5.88] | 1.30 [0.00, 10.29] | 3.10 [0.00, 6.58] | 0.673 | 0.828 |
| **PP (median [IQR])** | **17.73 [10.06, 34.11]** | **12.67 [6.88, 24.62]** | **8.49 [3.86, 14.04]** | **0.012** | **0.036** |
| SE (median [IQR]) | 5.85 [1.98, 13.01] | 5.59 [1.79, 10.31] | 3.16 [0.00, 13.16] | 0.458 | 0.692 |
| TB (median [IQR]) | 0.00 [0.00, 5.97] | 0.00 [0.00, 4.31] | 0.64 [0.00, 10.28] | 0.782 | 0.918 |
